# Glycosphingolipids Regulate Phosphatidylserine Transport at ER–PM Contact Sites

**DOI:** 10.1101/2025.09.26.678863

**Authors:** Ritchel Gannaban, Sher Ali, Wei Chen, Melanie Nguyen, Travis I. Moore, Yubin Zhou, John F. Hancock, Junchen Liu

**Author notes:** Corresponding authors: Junchen Liu,; John F. Hancock,; Yubin Zhou,.

## Abstract

Plasma membrane (PM) localization of KRAS requires specific glycosphingolipids in the outer leaflet and phosphatidylserine (PS) in the inner leaflet. PM PS content is controlled by lipid transport proteins ORP5 and ORP8, which operate at ER-PM membrane contact sites (MCSs). Using high-resolution imaging, we now show that GSLs including GM3 and SM4, are required to maintain ORP5 and ORP8 localization to MCSs. Genetic deletion or pharmacologic inhibition of enzymes required for the biosynthesis of GM3 or SM4, displace PI4-kinase Type IIIα (PI4KIIIα) and its adaptor EFR3A from the PM, thereby reducing PM phosphatidylinositol 4-phosphate (PI4P) content. PM interactions of ORP5 and ORP8 are also disrupted. Since ORP5 and ORP8 transport PS to the PM by counter-transporting PI4P to the ER, PM PS content is substantially reduced. We further show that GM3 and GM2 regulate the assembly of ER-PM-MCSs, such that the function of other MCS-localized macromolecular machineries including calcium release-activated calcium channels is abrogated when glycosphingolipid biosynthesis is blocked. Together, this study establishes glycosphingolipids as organizers of PS transport and ER-PM MCSs, expanding the regulators of MCSs beyond protein tethers to include glycosylated lipids and revealing how glycosphingolipids control KRAS function.

## Introduction

KRAS is a small GTPase that regulates a plethora of cellular processes including cell proliferation and differentiation. It operates as a bimodular switch that oscillates between a GDP-bound inactive form and a GTP-bound active form. Gain of function mutations at G12, G13 or Q61 frequently occur in pancreatic and lung cancer, which permanently lock KRAS in its GTP bound state, leading to its constitutive activation. For both mutant and wildtype KRAS to signal, plasma membrane localization and formation of nanoclusters are required^1–3^. These nanoclusters serve as hubs for KRAS interactions with effectors and regulatory proteins, which permit high-fidelity and low noise signal transduction^4–6^. PM association of KRAS is determined by its C-terminal membrane anchor that comprises a farnesyl cysteine and the adjacent polybasic domain (PBD) of six continuous lysines^7, 8^. The precise amino acid sequence of the prenyl group and PBD encodes a unique lipid preference of KRAS for unsaturated mixed chain phosphatidylserine (PS)^9, 10^. Depletion of PS from the PM, either through genetic approaches or pharmacological agent fendiline, result in loss of PM KRAS and impaired KRAS functions^10–13^. Beyond PS, we recently demonstrated that a subset of glycosylated PM lipids functions as high-level regulators of KRAS signaling. Glycosphingolipids (GSLs), exclusively expressed on the cell surface, comprise diverse group of molecules with a ceramide backbone linked to distinct sugar moieties^14^. Aberrant expression of GSLs has been implicated in multiple diseases including cancer, and neurological disorders^15^. We recently discovered that specific GSLs including GM3 (monosialodihexosylganglioside) and SM4 (3-O-sulfogalactosylceramide) are upregulated in KRAS mutant pancreatic cancer. Genetic deletion of UGCG, which encodes UDP-glucose ceramide glucosyltransferase, the first enzyme in GSL biosynthesis, reduces KRAS plasma membrane (PM) localization and nanoclustering, thereby impairing KRAS-driven oncogenesis. Concordantly, GSL depletion results in loss of PM PS, which are required for KRAS PM interaction and clustering^16^. However, the detailed mechanism underlying this process remains elusive.

Transport of PS between ER and PM by the lipid transport proteins ORP5 and ORP8 (oxysterol-binding protein related protein 5 and 8), is a key mechanism that maintains PS homeostasis at the PM^17^. ORP5 and ORP8 possess C-termini that anchor them to the ER and pleckstrin homology (PH) domains that interact with PI4P at the PM. The functional lipid transfer (ORD) domain then transports one PS from the ER to PM at the expense of the counter transport of a PM PI4P to the ER. Sac1, at the ER, then hydrolyzes PI4P to PI, producing a PI4P gradient at the MCSs^18^. The cycle is completed by delivery of PI back to the PM whereby it is phosphorylated by PI4KIIIα to regenerate PI4P^19^. Thus, efficient PS transport by OPR5 and ORP8 is dependent on the PI4P levels at the PM, and by inference, the activity of the PI4KIIIα complex^17^. Key components of the complex include TTC7 and FAM126A, which stabilize the PI4KIIIα at the inner leaflet PM surface, and EFR3A, an adaptor protein that anchors the complex to the PM through its palmitoylated cysteines at the N-terminus^20, 21^. Importantly, ORP5 and OPR8 tightly tether ER-PM membranes to form distinct MCSs that are conducive for lipid transport^17^. This PS transport machinery is crucial for KRAS function. Deletion of ORP5 and/or ORP8 cause loss of PS and KRAS from the PM, and reduced KRAS-dependent cell growth and oncogenesis^11, 12^. The PI4KIIIα signaling complex is also implicated in regulation of PM KRAS, as well as KRAS tumorigenicity^11, 13^. In this study, we combine high-resolution imaging, genetic manipulation, and patch-clamp electrophysiology to demonstrate that outer leaflet GSLs regulate inner leaflet PS content by precisely modulating key components of the PS transport machinery and associated ER–PM membrane contact sites.

## Results

### Depleting GSLs reduced PM PS content by disrupting assembly of ORP5 and ORP8

GSL depletion abrogates KRAS PM interaction and clustering by reducing inner leaflet PS content. This can be quantified in MDCK cells expressing GFP-LactC2, a genetically encoded biosensor that possesses a PH domain which specifically binds to the headgroups of PS. PM sheets prepared from these cells are labeled with anti-GFP antibodies coupled to 4.5nm gold particles and visualized by electron microscopy (EM). Treatment with DL-PDMP, an inhibitor of UDP-glucose ceramide glucosyl transferase (UGCG), the first enzyme in ganglioside biosynthesis, significantly reduces PM PS content as quantified by Lact-C2 gold labeling intensity (per μm^2^) compared with control cells (Fig.1A). Lipid addback experiments in these cells showed that 1hr incubation with GM3 (monosialodihexosylganglioside), a simple ganglioside fully restored the PM PS content (Fig. 1A), whereas a 1hr incubation with SM4 (3-O-sulfogalactosylceramide), a simple sulfatide, partially restored PM PS levels. To explore the mechanism underlying this reduction in inner leaflet PM PS we first conducted shot gun lipidomics in cells depleted of GSLs by UGCG-inhibition, or glucose starvation^16^. We observed an increase in PS levels after UGCG inhibition (Fig.1B). This result indicates that reduced PM PS content upon GSL depletion is not a consequence of a reduction in total cellular PS levels.

**Fig. 1.**
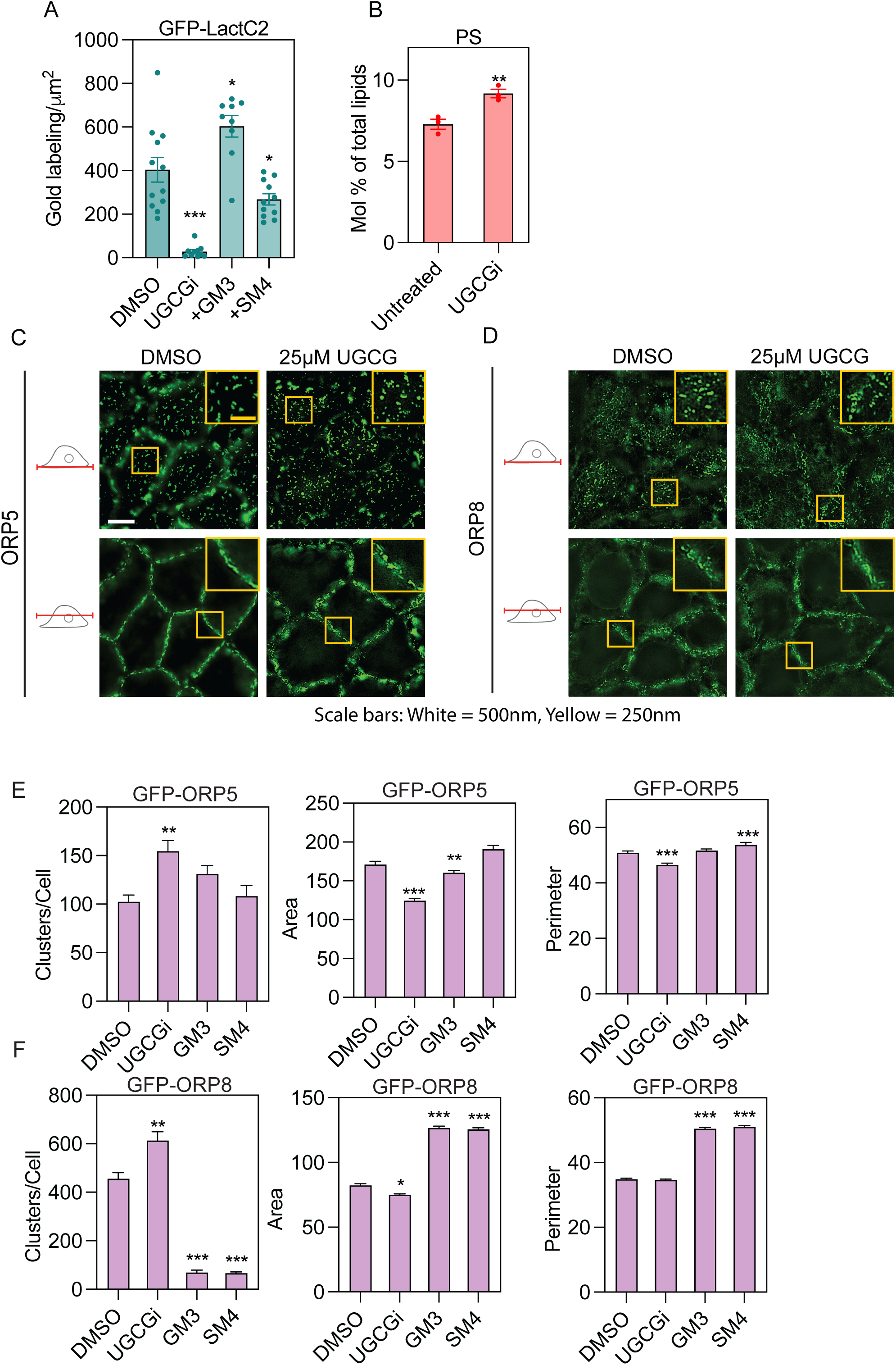
Suppressing GSL biosynthesis perturbs PS transport mediated by ORP5/8. **(A)** MDCK cells stably expressing GFP-LactC2 (PS probe) treated with vehicle (DMSO) or 25μM DL-PDMP (UGCG inhibitor) (24h) and incubated with exogenous GM3 or SM4 for 1hr were evaluated using EM. PM GFP-LactC2 labeling were quantified as the mean gold labeling intensity (± SEM, n ≥ 12). **(B)** Molar fraction of PS headgroup from the lipidomics analysis of MDCK cells treated with vehicle or UGCG inhibitors (n = 3, ± SD). Two tailed student’s t-tests were used to evaluate significant differences in gold densities and molar fractions compared to control cells (*p < 0.05, **p < 0.01, ***p < 0.001). **(C-F)** SIM super-resolution imaging of MDCK cells stably expressing GFP-ORP5 **(C)** or GFP-ORP8 **(D)** and treated with DMSO or 25μM DL-PDMP for 24h. **(C-D)** Representative images of basal membrane and cross sections of the cells. Rectangular regions enclosed by yellow boxes are highlighted in insets. Scale bars: White = 500nm, Yellow = 250nm. **(E-F)** Quantification and statistical analysis of basal membrane GFP-ORP5 or ORP8 cluster segmentation from MDCK cells stably expressing GFP-ORP5 or ORP8 and treated with vehicle or a UGCG inhibitor for 24h with or without exogenous GM3 or SM4 addback for 1h prior to SIM imaging. Cluster number, area, and perimeter of ORP5 or ORP8 clusters were quantified using CellProfiler 4.0. (± SEM n ≥ 6). Differences from control were evaluated in two-tailed student’s t-tests (*p < 0.05, **p < 0.01, ***p < 0.001).

PS transport from the ER to the PM, is mediated by ORP5 and ORP8, which have been shown previously to be required for KRAS PM targeting. We therefore examined whether UGCG inhibitors affect the subcellular distributions of ORP5 and ORP8. Structured illumination microscopy (SIM) super-resolution z-stacks were acquired from the basal PM through the cellular cross section of MDCK cells expressing GFP-ORP5 or GFP-ORP8. GFP positive clusters were subsequently segmented, and their intensity and morphological parameters quantified. Consistent with previous reports^17, 22^, GFP-ORP5 formed clearly defined puncta at the cell periphery reflecting localization to ER-PM MCSs, whereas GFP-ORP8 puncta were visualized more widely on both the PM and morphological ER (Fig. 1C and D). Overnight treatment with DL-PDMP led to an increased number of smaller GFP-ORP5-decorated puncta, as indicated by a higher cluster count per cell and reduced cluster area and cluster perimeter (Fig. 1C). To test GSL specificity of the effect, we incubated DL-PDMP treated cells with GM3, a downstream product of GlcCer, into UGCG-depleted cells. GM3 addback reduced the GFP positive puncta count while increasing cluster area and perimeter, effectively reversing the effects of UGCG inhibition. Similarly, adding back SM4, a GalCer series GSL, also reversed the changes in GFP-ORP5 puncta number and morphology (Fig. 1C and E, and s1). We repeated these experiments in MDCK cells expressing GFP-ORP8. DL-PDMP treatment increased the number of GFP-positive ORP8 puncta while reducing their area. Interestingly, compared to baseline levels (DMSO), 1hr incubation with GM3 and SM4 led to a 45% increase in cluster area, a 54% increase in perimeter, and an 85% reduction in puncta count (Fig. 1D and F, and S1), indicating that GM3 and SM4 have a stronger impact on the number and size of GFP-ORP8 puncta than GFP-ORP5 puncta. Overall, these results strongly indicate that the glycosphingolipids GM3 and SM4 modulate the number and size of the MCSs containing ORP5 and ORP8.

### GSLs are required for the PM interactions of ORP5 and ORP8

We next directly quantified the PM interactions and clustering of ORP5/8 using univariate EM analysis. PM sheets of Caco-2 cells expressing GFP-ORP5 or GFP-ORP8 with CRISPR deletion of UGCG expression, were attached to EM grinds, fixed, labelled with 4.5nm gold-anti-GFP, and imaged by EM. Analysis of the gold distribution patterns revealed that UGCG knockout mislocalized ORP5 from the PM, evident by a reduction in gold labeling intensity, but with no effect on the clustering of GFP-ORP5 that remained attached to the PM, as quantified by the max value (*L_max_*) of the gold pattern K function expressed as *L(r)-r* (Fig. 2B and s2A).Lipid addback experiments showed that addback of GM3, but not SM4, restored the PM interaction of ORP5 (Fig. 2A). Similarly, UGCG deletion significantly mislocalized GFP-ORP8 from the PM in Caco-2 cells, which was partially rescued by GM3 and SM4 (Fig. 2C and D). Overall, these findings demonstrate that biosynthesis of specific GSLs is essential for the PM targeting of ORP5 and ORP8.

**Fig 2.**
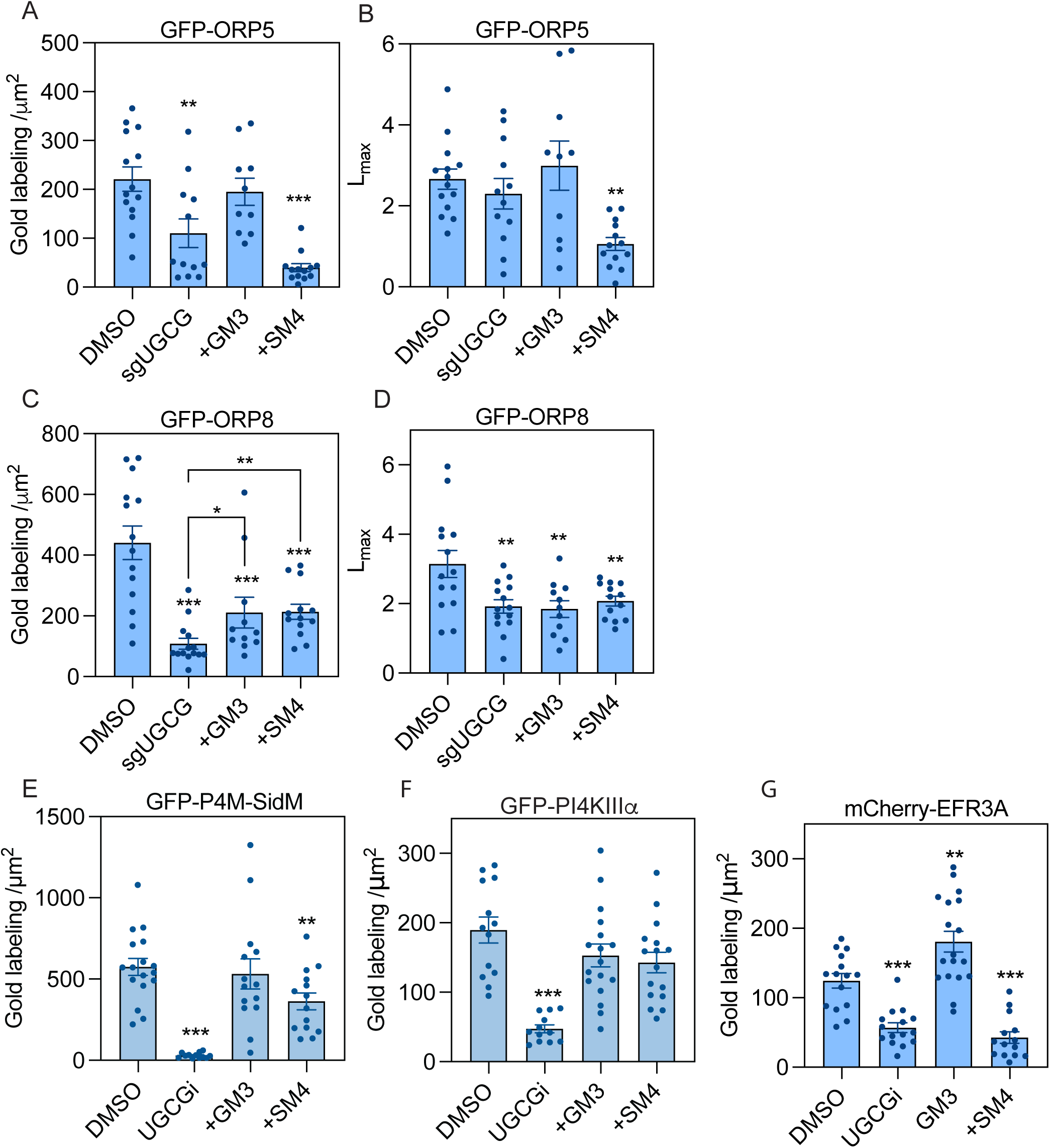
GSLs are required to maintain PM localization of ORP5/8, the PI4KIIIa complex, and PM PI4P levels. **(A-D)** Caco-2 cells transiently expressing GFP-ORP5 or GFP-ORP8 were depleted of UGCG using CRISPR-cas9 (sgUGCG) and compared with non-targeting control (sgCtrl). Cells were incubated with exogenous GM3 or SM4 for 1h before PM sheets were prepared, immunogold labeled with 4.5nm-gold-anti-GFP, and imaged by EM. PM interactions of ORP5 **(A)** and ORP8 **(C)** were quantified as mean gold labeling density (± SEM; n ≥ 15 for each condition). The extents of nanoclustering for ORP5 **(B)** and ORP8 **(D)** were quantified as the maximum value (*L_max_*, ± SEM; n ≥ 15) of the *L(r)-r* function. **(E-G)** MDCK cells transiently transfected with GFP-P4M-SidM **(E)**, GFP-PI4KIIIa **(F)**, or mCherry-EFR3A **(G)** were incubated with a UGCG inhibitor (DL-PDMP) for 24h and subsequently incubated with GM3, or Gb3 for 1 h. PM sheets of the cells were labeled with 2nm-gold-anti-mCherry or 4.5nm-gold-anti-GFP antibodies and visualized by EM. The PM abundance of PI4P, PI4KIIIa, EFR3A, and ER-PM contact sites were quantified as gold labeling density. (B and D) Significant differences between *L_max_*values for GSL-addback and control cells were evaluated in bootstrap tests (*p < 0.05, **p< 0.01). (A, C, E-G) Statistical differences from control were evaluated using t-tests (± SEM, n ≥ 15, *p < 0.05, **p < 0.01, ***p < 0.001). GFP-P4M-SidM: PIP4 probe.

### Suppressing GSL biosynthesis displaced the PI4KIII**α** complex from the PM

ORP5 and ORP8 are anchored to the PM through interaction between a PH domain and PI4P^17^. We therefore quantified PM PI4P levels in GSL-depleted cells using immunogold labeling of a PI4P biosensor, GFP-P4M-SidM. EM analysis showed that DL-PDMP treatment markedly reduced PM levels of PI4P. GM3 addback fully recovered PI4P levels, whereas SM4 addback partially rescued PI4P levels (Fig. 2E). PI4P is synthesized by PI4KIIIα so to explore further, we quantified the PM levels of PI4KIIIα, which localizes to the PM by a mutlitiprotein complex that includes an anchoring protein EFR3A. EM analyses in MDCK cells transiently expressing GFP-PI4KIIIα or mCherry-EFR3A showed that UGCG inhibition mislocalized both proteins. GM3 addback efficiently restored the PM localization of PI4KIIIα and EFR3A (Fig. 2F and G), whereas addback of SM4 fully recovered PI4KIIIα to the PM but had no effect on EFR3A, again reflecting differential roles of these two GSLs (Fig. 2F and G). Taken together, these data demonstrate that GM3 and SM4 control PM localization of ORP5/8 by modulating PM localization of the PI4KIIIα complex.

### Endogenous GM3 and SM4 are required for the correct assembly of PS transport machinery

Lipid addback results suggest that GM3 and SM4 play critical but distinct roles in regulating different components of the PS transport machinery. To investigate this further, we deleted specific enzymes involved in GM3 or SM4 biosynthesis, including *ST3GAL5*, which encodes GM3 synthase (GM3S) responsible for converting LacCer to GM3, and *UGT8*, which encodes UDP-galactose ceramide galactosyltransferase, the enzyme that synthesizes GalCer, the immediate precursor of SM4. Deletions of either *ST3GAL5* or *UGT8* significantly disrupted the PM localization and clustering of ORP5, which was fully recovered by addback of cognate missing GSL (Fig. 3A and B, and s2B and C). Likewise, we observed a loss of PM-localized ORP8 and diminished ORP8 clusters in cells depleted of GM3 or SM4 by deletion of GM3S or UGT8 respectively. In the knockout cells, GM3 or SM4 addback fully recovered the PM targeting and clustering of ORP8 (Fig. 3C and D).

**Fig 3.**
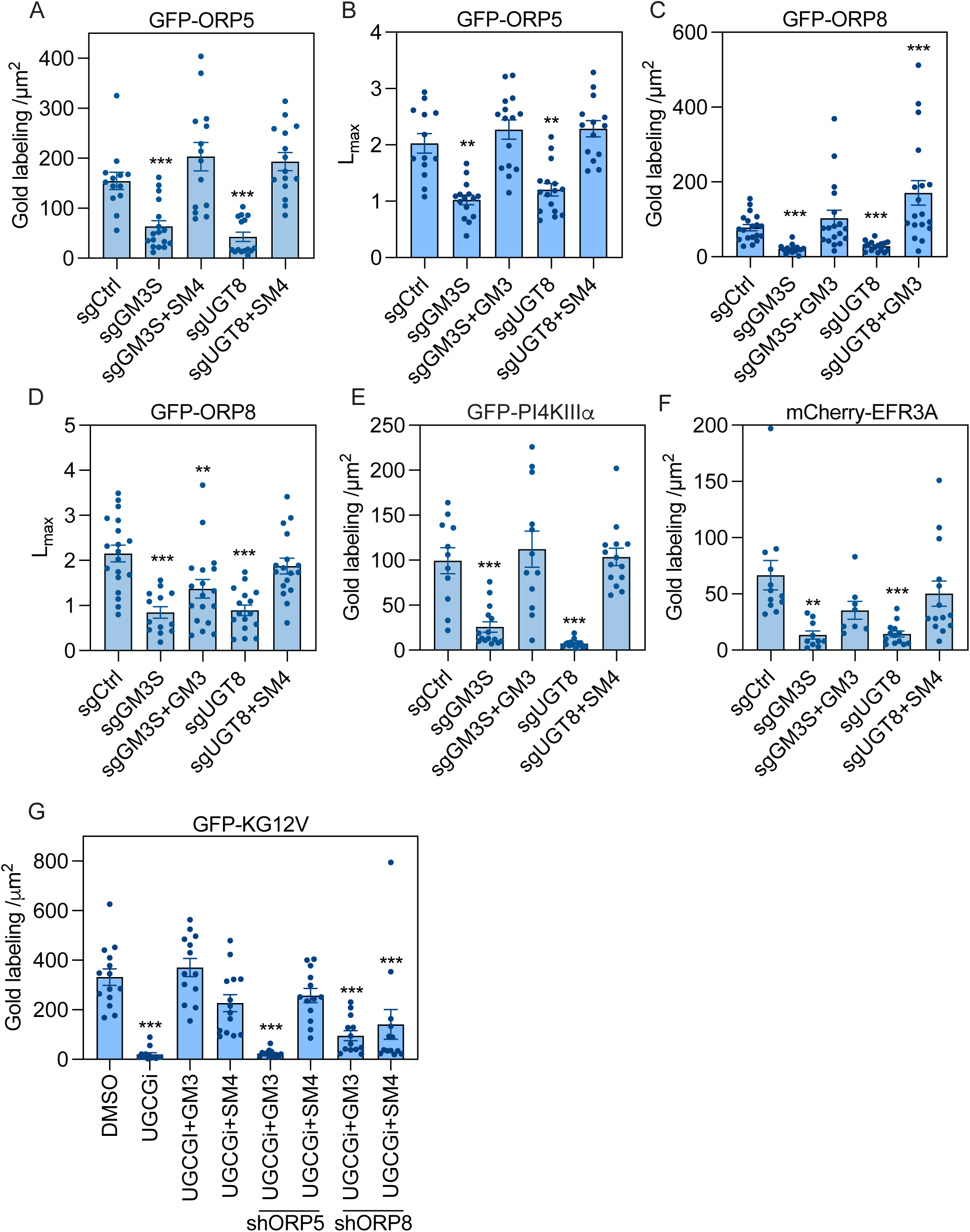
Key biosynthetic enzymes for GM3 or SM4 are required for PS transport machinery. **(A-F)** Caco-2 cells depleted of *ST3GAL5* (sgGM3S) or *UGT8* (sgUGT8), or expressing control sgRNAs (sgCtrl), were transiently transfected with GFP-ORP5, -ORP8, -PI4KA, or mCherry-EFR3A. GM3 or SM4 was then reintroduced to *ST3GAL5* or *UGT8* knockout cells, respectively, before EM analysis of PM sheets. PM localization of GFP-ORP5 **(A)**, -ORP8 **(C)**, -PI4KIIIα **(E)** or mCherry-EFR3A **(F)** was quantified by gold labeling density. Extent of nanoclustering of ORP5 **(B)** and ORP8 **(C)** was quantified by the maximum value (*L_max_*) of *L(r)-r*. **(G)** Caco-2 cells stably expressing GFP-KRASG12V with or without ORP5 or ORP8 knockdown were treated with DMSO or DL-PDMP for 24h. The cells were then incubated with GM3 or SM4 for 1 before fixed for EM analysis. PM localization of GFP-KG12V were quantified as gold labeling density. (**A**, **C**, and **E-F**) Statistical differences from control were evaluated using t-tests (± SEM, n ≥ 12, *p < 0.05, **p < 0.01, ***p < 0.001). (**B** and **D**) Bootstrap tests were used to evaluate significances in Lmax (± SEM, n ≥ 12, *p < 0.05, **p< 0.01)

We next examined the impact of genetic deletion of these enzymes on the PI4KIIIα complex. We found that deletion of GM3S or UGT8 significantly reduced the levels of PM-localized PI4KIIIα and its membrane anchor EFR3A. Full recovery of PI4KIIIα and EFR3Ato the PM was observed upon addback of GM3 or SM4 into the GM3S and UGT8 knockout cells, respectively (Fig. 3E and F). Taken together, these results demonstrate endogenous GM3 and SM4 are required for the PM localization of key components of PS transport machinery.

To assess the relevance of these findings in KRAS mutant cells, we assessed the ability of GM3 or SM4 addback to restore PM KRASG12V levels in ORP5 and ORP8 knockdown cells. Consistent with our previous report, UGCG inhibitors significantly reduced the PM localization and nanoclustering of KRASG12V in Caco-2 cells (Fig. 3G). GM3 addback fully restored KRAS PM localization in UGCG inhibited control cells but was not able to restore KRAS in either UGCG inhibited ORP5 nor ORP8 knockdown cells. By contrast SM4 addback partially restored PM localization of KRAS in both ORP5 deleted and control cells, but not in ORP8 knockdown cells.

### Formation of ER-PM MCSs are contingent on GSLs

ORP5 and ORP8 operate as PS transporters at ER-PM contact sites, where the two membranes are closely apposed (∼10–30 nm) to facilitate efficient non-vesicular lipid transport^18^. To quantitatively analyze if GSLs might regulate the assembly of the MCS, we directly visualized distribution of the MCS probe GFP-Mapper^23^ using EM. The GFP moiety of GFP-Mapper is expressed on the luminal side of the ER such that on removal of the apical PM together with tightly opposed ER membrane onto an EM grid, ER-localized GFP becomes solvent exposed. Thus, GFP-Mapper at the cortical ER can be labeled by 4.5 nm gold-conjugated anti-GFP, allowing nanoscale visualization of ER-PM contact sites by EM imaging (Fig. 4A). This novel approach may provide higher resolution and a more quantitative method for analyzing ER-PM contact sites than conventional confocal imaging. In this assay we observed a strong immunogold labeling of GFP-Mapper in a clustered distribution which intriguingly was significantly reduced by treatment of DL-PDMP (Fig. 4B), suggesting that GSLs are required for the assembly of ER-PM contact sites. Next, we incubated DL-PDMP treated cells with defined GSLs for 1h before preparation of the PM sheets. While GM3 addback partially restored ER-PM contact sites, GM2, which contains an additional N-acetylgalactosamine, fully restored GFP-Mapper labeling. In contrast, SM4 had no effect on the ER-PM contact sites labeled by GFP-mapper (Fig. 4B and C). These data, strongly suggest that ER-PM contact sites are regulated by the gangliosides GM3 and GM2.

**Fig 4.**
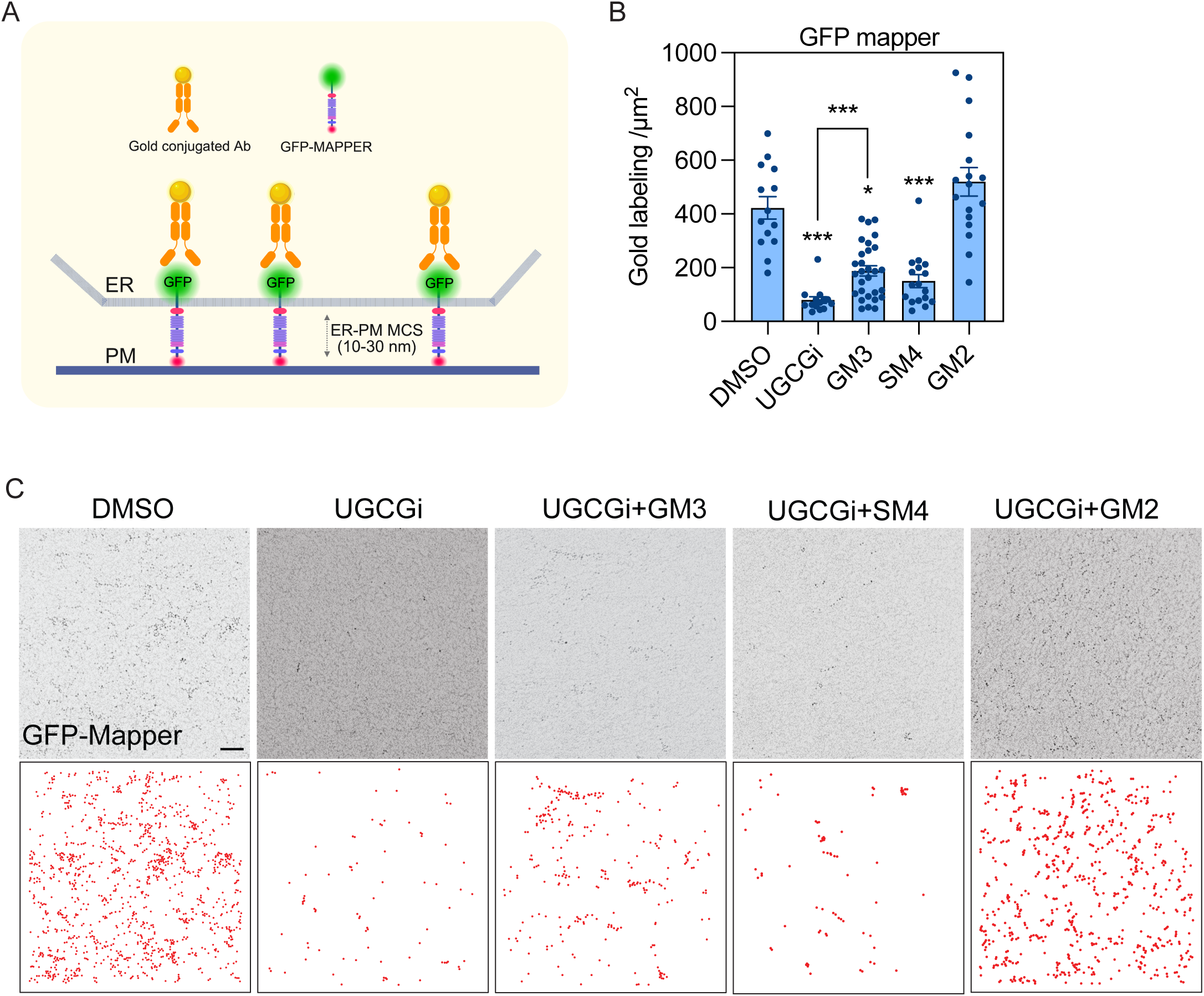
Formation of ER-PM contact sites are dependent on specific GSLs. **(A)** Schematic illustration of labeling the ER-PM contact site marker GFP-Mapper using 4.5 nm gold conjugated anti-GFP antibodies. **(B-C)** MDCK cells stably expressing GFP-Mapper were treated with DL-PDMP for 24hr and incubated with GM3, GM2, SM4 and Gb2 for 1hr. Cortical ER membrane sheets apposed to the basal PM were attached to EM grids. Removal of the apical PM exposed the GFP tag expressed on the cytosolic side of the ER. The samples were then fixed and immunolabeled with anti-GFP antibodies conjugated to 4.5 nm gold particles, enabling effective quantification of ER-PM contact site abundance based on gold labeling intensity. (**A**) Representative and digitalized images of ER-PM contact sites that are labeled by its marker GFP-Mapper at 100 x magnification. Scale bar: 100nm. (**B**) The abundance of the ER-PM contact sites are quantified the gold labeling density. Statistical significance relative to the control was assessed by two-tailed Student’s t-tests (± SEM, n ≥ 15, *p < 0.05).

### Diverse functions of ER-PM contact sites are regulated by glycosphingolipids

The ER-anchoring component of GFP-Mapper is derived from stromal interaction molecule 1 (STIM1), a Ca²⁺ sensor in the endoplasmic reticulum (ER). Upon depletion of ER Ca2+, STIM1 undergoes conformational change and translocates to ER-PM MCSs to gate ORAI1, a pore-subunit in the PM. Together STIM1 and ORAI1 constitute the Ca2+ release-activated Ca2+ (CRAC) channel. Given the requirement for GSLs to maintain the PM interactions of GFP-Mapper we next examined localization of full-length STIM1 at ER-PM MCSs and operational integrity of CRAC channels. To assess this, PM sheets from HeLa cells expressing STIM1-mCherry were fixed and labeled with 6nm-gold-anti-mCherry antibodies for EM imaging. Following the similar logic in Fig 3A, since only PM-tethered STIM1 can be detected by EM, this approach provides a quantitative assessment of STIM1 at MCSs. Statistical analysis of gold particle distribution revealed that DL-PDMP treatment significantly reduced STIM1 localization and clustering at the PM (Fig. 5A). We next conducted lipid addback experiments under DL-PDMP treatment. Exogenous SM4 partically rescued STIM1 attachment to the PM, while GM3 fully restored it (Fig. 5A).

**Fig 5.**
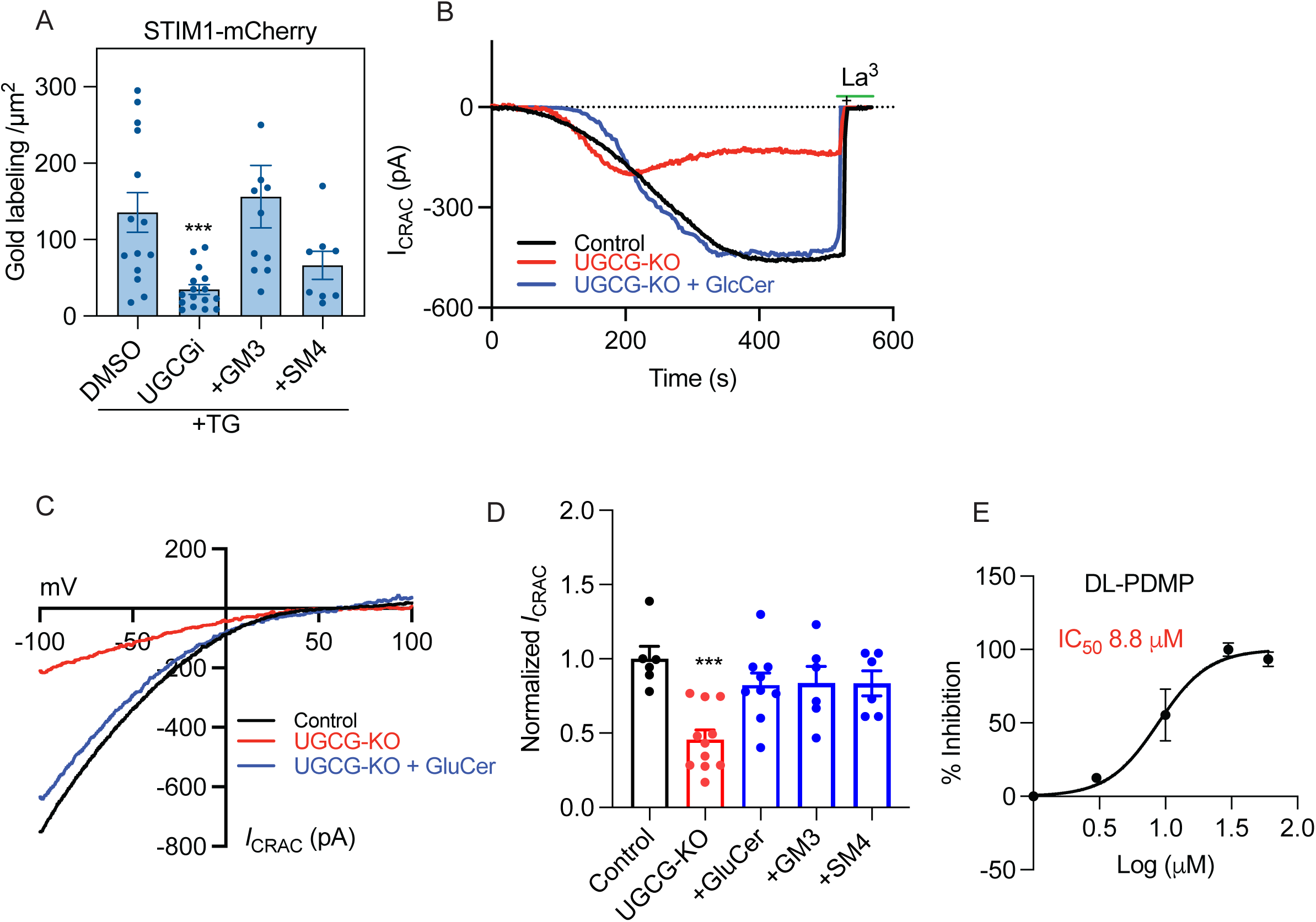
Glycosphingolipid inhibition suppresses calcium signaling at the ER-PM contact sites. **(A)** EM evaluation of PM binding of STIM1 in cells treated with DL-PDMP and incubated with GlcCer, GM3 or SM4, and thapsigargin (TG) was added to activate the STIM1-ORAI1 channel. (**B)** Representative current-time relationship of whole cell CRAC current density (*I_CRAC_*) evaluated at −100 mV activated upon passive store-depletion of HeLa cells (control, UGCG-KO and Pre-treated UGCG-KO with GlucCer for 3-4 hours) stably transfected Orai1-GFP with transiently expressed STIM1-mCh. La^3+^ was applied to subtract the leak current. Compared with control, UGCG-KO HeLa cells displayed significantly decreased *I_CRAC_*, and was restored with the pre-treamtment with GluCer. (**C**) Representative I-V relationship corresponding to the peak current observed in (B). **(D)** The bar graph summarizes the normalized current densities (mean±SEM) in control (n=10), UGCG-KO (n=11) and pre-treated with GlucCer, GM2, GM3 and SM4 (n=6 to 9 each). (*** *P* value < 0.001, unpaired student’s t-test). (**E)** UGCG inhibitor DL-PDMP blocks CRAC channel with an IC_50_ of 8.8 µM.

**Fig 6.**
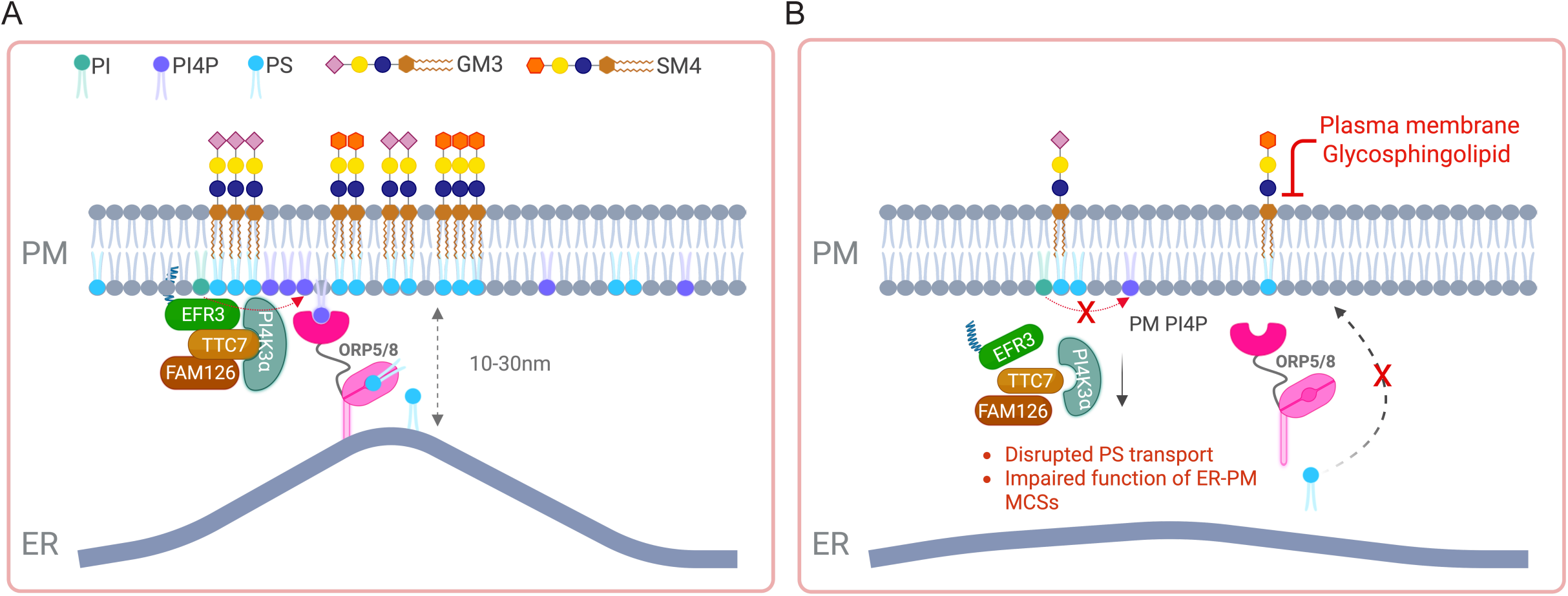
Schematic of how GSLs regulate the PS transport machinery and associated MCSs. **(A)** GSLs are required for PM anchoring of the PI4KIIIα complex, which produces PI4P at the PM. PM PI4P is transported to the ER by ORP5 and ORP8 in exchange for PS delivery from the ER to the PM. Successful lipid transport depends on the formation of ER–PM MCSs, mediated by tethers such as ORP5/8 and the STIM1–ORAI1 complex. **(B)** In the absence of outer leaflet GSLs, the PI4KIIIα complex is displaced from the PM, and the attendant PS transport is abolished due to the loss of PM PI4P. GSL deletion disrupts the assembly and function of ER– PM MCSs. Taken together, GSLs act as high-level regulators that orchestrate PS transport machinery at ER–PM MCSs.

Finally, we conducted patch clamping on HEK293 cells stably expressing STIM1/ORAI1, with and without UGCG depletion. We found that UGCG depletion significantly reduced the CRAC current (I_CRAC_; Fig. 5B and C). The suppressive effect was rescued by add-back of GlcCer, GM3 or SM4 in the culture media (Fig. 5D and E). Concordantly, UGCG inhibitors potently suppressed I_CRAC_ in a dose dependent manner (Fig. 5E) These data, along with the results from the EM analyses, strongly indicate that GSLs act as high-level regulators of the assembly and function of ER-PM contact sites.

## Discussion

Here we show that GM3 and SM4 are directly involved in maintaining the integrity of the PS transport machinery at the ER-PM contact sites. Several lines of evidence support this important conlusion. First, we observed mislocalization of ORP5 and ORP8 in cells treated with a UGCG inhibitor. UGCG inhibition also caused loss of PI4KIIIα and its anchor protein EFR3A from the PM, which resulted in a substantial decrease in PM PI4P levels. PI4P together with phosphatidylinositol 4,5-bisphosphate (PIP_2_) are required to maintain the localization of ORP5 and ORP8 to PM-ER MCSs, explaining the mislocalization of ORP5 and ORP8 when GSLs were depleted. Secondly, acute addback of GM3, or SM4 fully or partially recovered PM PI4P respectively, and restored PM localization of EFR3A and PI4KIIIα. Similar effects on PM PI4P with the same attendant consequences on ORP5/8 localization and reduced PM PtdSer levels occur in cells treated with specific PI4KIIIα inhibitors ^12, 13^. Thirdly, the role of GSLs likely extend beyond simple maintenance of PI4KIIIα PM localization because ER-PM MCSs *per se* disassembled upon UGCG inhibition and were partially or fully restored by addback of GM3 or GM2, respectively, strongly suggesting an additional structural or operational role for theses GSLs in ER-PM MCS integrity. How such a role might integrate with known membrane tethers such as VAMP-associated proteins and extended synaptotagmins ^24–26^ remains to be explored. Moreover, the emerging role of PI4P in defining PM identity ^27, 28^ by contributing to the synthesis of PM polyanionic lipids, and lipid counter exchange across LTPs to transport PtdSer to the PM, by extension now calls for a similar defining role for the GSLs GM3/2 and SM4. At a molecular level this will likely involve the biophysical properties of GSL long acyl chains organizing or generating lipid-based platforms, or domains ^29–31^, that facilitate assembly of the multiprotein complex required for PI4P generation. Operationally this may extend to the PI-LTPs: PITPα/β, PITPNC1, PITPNM1 and PITPNM2 ^32^, which support ORP5 and OPR8 for PtdSer delivery, as well as defining PM sites for the binding of ER-PM MCS structural proteins.

More broadly the study reveals functional coupling of two distinct metabolic pathways: GSL biosynthesis and non-vesicular PS transport. Supporting this notion, a recent study showed that the heterodimerization of ORP9 and ORP11, LTPs that exchange PS and PI4P at the ER-trans-Golgi contact sites, is required for sphingomyelin synthesis. Knockout of ORP9 or ORP11 resulted in reduced sphingomyelin levels in Golgi apparatus^33^. Although the authors did not evaluate whether manipulating sphingomyelin metabolism also affects PS/PI4P exchange at the MCSs, their findings, together with ours provide strong evidence that sphingolipid metabolism is tightly linked to non-vesicular trafficking of PS across different membranes. Given these observations, it will be interesting to explore whether ORP5 and ORP8 reciprocally impact trafficking or biosynthesis of GSLs including GM3 and SM4.

Our findings that GSL depletion suppresses CRAC channel activity extend the lipid regulators of STIM1 to outer leaflet GSLs. Initial studies on CRAC channel predominantly examined PIP_2_, a negatively charged phospholipid that interacts with the polybasic domain (PB) at the C-terminus of STIM1^34^. Cohen et al. recently expanded the known phosphoinositide-binding region to include the SOAR (STIM-ORAI activation region) of STIM1. Their study identified a critical lysine stretch (^382^KIKKK^386^) within the first helix of SOAR, which is essential for STIM1 clustering at ER-PM junctions following Ca²⁺ store depletion^35^. Mutating these lysines to alanines (STIM1-4KA) completely disrupted ER-PM MCS formation. Further biochemical and biophysical analyses demonstrated that SOAR binding to PI4P plays a crucial role in targeting STIM1 to the PM, as PI4P depletion markedly reduced STIM1 puncta formation at ER-PM junctions^35^. Given the critical role of PI4P in the MCSs defined by ORP5/8 and STIM1/ORAI1, the PI4KIIIα complex, whose PM localization we have now shown to be closely associated with GSLs, may play a key role in orchestrating MCS formation and its diverse functions. In this context, a recent study showed that manipulating the localization of ORP5 and ORP8 affects the clustering of STIM1 and STIM-CRAC current^36^, thereby linking PS/PI4P exchange to the store operated Ca^2+^ entry. Consistently, our findings here have shown that GSLs regulate both ORP5/8 and STIM1 at the ER-PM contact sites, suggesting that the functional integration of these two distinct protein complexes is contingent on PM lipids.

Previous work on MCSs has centered on the tether proteins that bridge two membranes through protein-protein or lipid-protein interactions. We now have expanded the regulators of PM-ER MCSs to include a class of outer leaflet sphingolipids. Interestingly, a disruption of MCS between ER and late endosomes and lysosomes ouccurs in cells depleted of sphingosine kinases 1 and 2 (SphK1 and 2), which convert sphingosine to S1P by phosphorylation^37^. SphK deletion reduces ER contacts with late endosomes and promotes PM recruitment of the cholesterol transporter Aster B, leading to an increase in PM cholesterol content^37^. Of note SphK deletion does not affect the subcellular localization of ORP5, which therefore appears to be primarily controlled by GSLs, rather than sphingosines or S1P, indication that glycosylation of ceramide represents a distinct metabolic fate with unique biological function.

In sum, our work demonstrates that outer leaflet GSLs act as high-level regulators of ER-PM contact sites, thereby controlling the functional assembly of key lipid transporters and Ca+ channels. Their cell surface localization may render them more accessible as drug targets than cytosolic proteins and lipids, which warrants future investigations.

