## Supplemental fig1-2 and Methods for "Glycosphingolipids Regulate Phosphatidylserine Transport at ER–PM Contact Sites"

Fig. s1

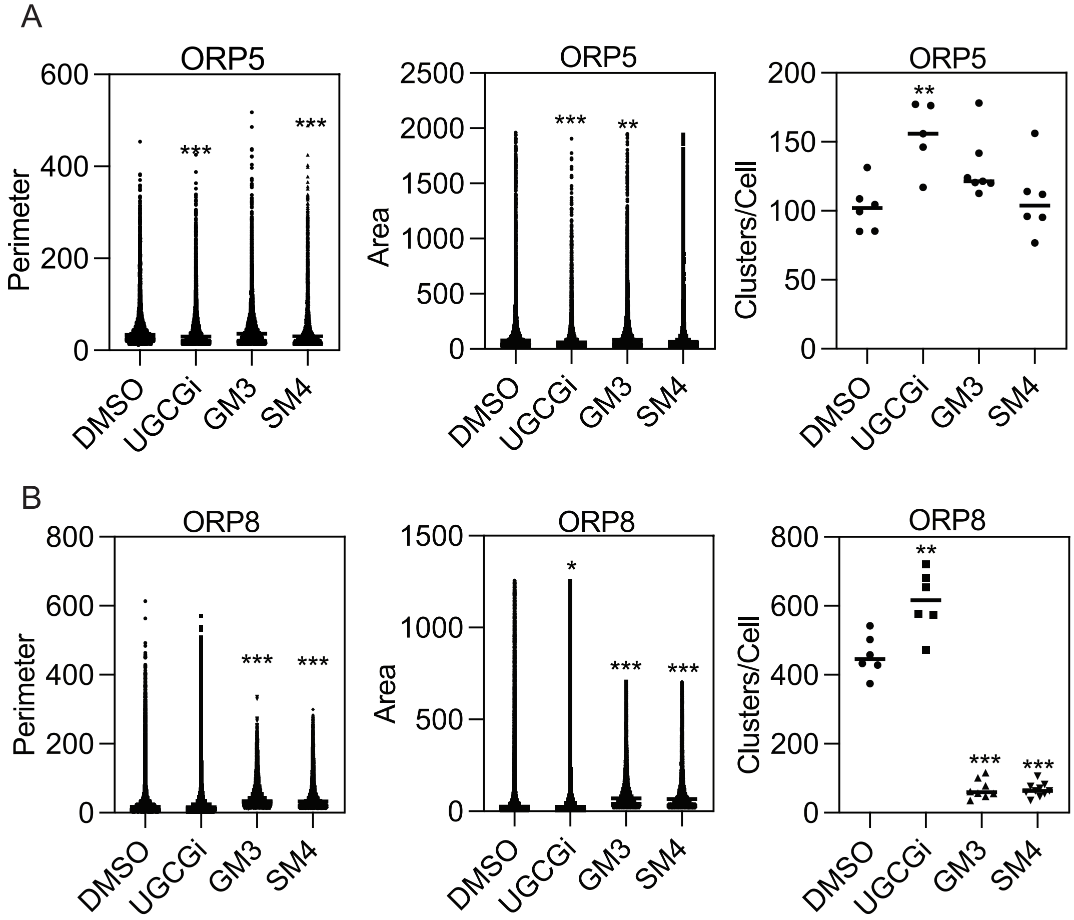

A
ORP5
Perimeter
600
400
200
0
***
***
DMSO
UGCGi
GM3
SM4
ORP5
Area
2500
2000
1500
1000
500
0
***
**
ORP5
Clusters/Cell
200
150
100
50
**
B
ORP8
Perimeter
800
600
***
***
ORP8
Area
1500
1000
*
ORP8
Clusters/Cell

Fig. s2

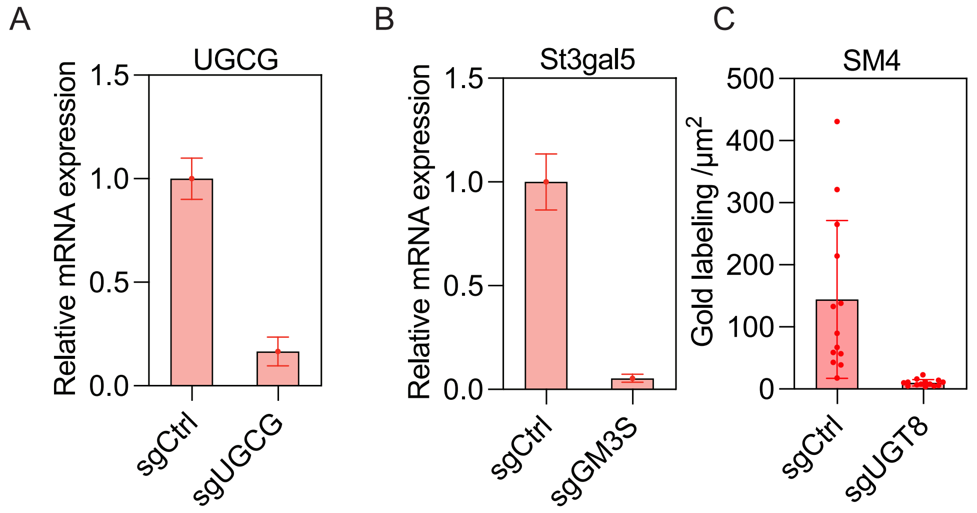
A
UGCG
Relative mRNA expression
1.5
1.0
0.5
0.0
sgCtrl
sgUGCG
B
St3gal5
Relative mRNA expression
1.5
1.0
0.5
0.0
sgCtrl
sgGM3S
C
SM4
Gold labeling /μm2
500
400
300
200
100
0
sgCtrl
sgUGT8

**Legends for supplementary figures**

**Fig s1. Suppressing sphingolipids altered the morphology of ORP5 and ORP8 puncta.**

(A-B) SIM super-resolution imaging of MDCK cells stably expressing GFP-ORP5 (A) or GFP-ORP8 (B) after 24-hour treatment with DMSO or 25 μM DL-PDMP. Graphs displaying individual data points for Fig. 1E and 1F are shown.

**Fig s2. Validation of knockout efficiency of selected GSL synthases.**

(A-B) qPCR analysis of relative mRNA levels of (A) UGCG or (B) st3gal5 (GM3 synthase) in Caco-2 cells expressing sgRNAs targeting UGCG or st3gal5, respectively. (C) EM evaluation of SM4 on plasma membrane sheets from Caco-2 cells deleted of UGT8 (sgUGT8), labeled with 4.5 nm gold-conjugated anti-SM4 antibodies.

### Materials and Methods

#### Materials

Dulbecco's modified eagle medium (DMEM) (Cat#11965092), glucose-free DMEM (Cat#11966025) and Fetal bovine serum (FBS) (Cat#A5670701) were purchased from GIBCO. Blasticidine S hydrochloride (Cat#15205) was purchased from Sigma and Puromycin (Cat#BP2956-100) was purchased from Thermo Fisher Scientific. The UGCG inhibitor DL-Threo-1-phenyl-2-decanoylamino-3-morpholino-1-propanol (DL-PDMP) was purchased from Enzo Biochem (#BML-SL210-0010). Lipofectamine™ 2000 transfection reagent (Cat # 11668030) was purchased from Thermo Fisher Scientific. Fixatives paraformaldehyde (PFA) and glutaraldehyde were available in house. The anti-GFP antibodies used for immunogold labeling were prepared in house. All lipids were purchased from Avanti Polar Lipids: Gb3 (#860699), GM3 (#860058), GlcCer (#860547), LacCer (#860590), and brain SM4 (#131305).

#### Lipid preparation

GlcCer, GalCer, GM3 and Gb3 were dissolved in chloroform/methanol mixture (2:1) while LacCer in choloform/methanol/water (5:1:0.1). The lipids are allowed to dry overnight by applying vacuum. On the day of the experiment, lipids were resuspended in a complete or glucose-free DMEM media with 10% FBS followed by sonication for 30 min. The suspension was kept from light and stored up to 4 h in 37°C. For the addback experiment, MDCK or Caco-2 cells were cultured in medium containing lipids for 1 h before EM analysis.

#### SIM imaging

The cells were imaged using structured illumination microscopy using a Nikon N-SIM super-resolution microscope system, equipped with a CIF App TIRF 100X objective (NA 1.49, Nikon Instruments) and images captured by an ORCA-Flash4.0 CMOS camera (Hamamatsu). Z-stacks were acquired every 0.2 μm over 10 μm and resulting 3D-SIM Z-stack dataset of 15 images per plane and reconstructed using the slice reconstruction method in NIS-Elements (Nikon Instruments). Images of the basal cellular surface and central cross-section were extracted from the slice and EGFP localization segmented using CellProfiler 4.0 (McQuin 2018) using the Otsu two class intensity thresholding method. The central cross-section localization was used to segment the cell membrane which was then projected onto the basal image to segment individual cells in the monolayer. Segmented EGFP clusters were then counted and area, perimeter, and mean intensity measured.

**Cell culture and transfection**

Mardin-Darby canine kidney (MDCK) cells gifted by Dr. Robert G Parton (University of Queensland, Australia) were maintained in DMEM supplemented with 10% FBS. All cells were cultured in 5% CO2 incubators at 37°C. For glucose starvation, MDCK cells were cultured in glucose-free DMEM for 4 hr. For immunofluorescence experiments, cells were seeded in six-well plates containing two gold grids (product cat and company) per well at a density of 2 x 105 cells/well 1 day prior to experiment. For transfection, MDCK cells were seeded at a density of 4 X 105 in 3.5 cm dish in growth medium without antibiotics 1 day before transfection. The next day, cells were transfected with plasmid expressing either GFP-KRASG12V, ORP5, or ORP8 with pEF6 promoter using lipofectamine 2000. On day 3, antibiotic selection was done with blasticidin at 10μg/ml. Cells were passaged 3 times before seeding in 96-well plate to generate single cell lines exhibiting optimal and uniform GFP-KRASG12V or GFP-OPR5 or GFP-ORP8 expressions.

**Generation of CRISPR-Cas9 cell lines**

The sgRNAs targeting human UGCG (5′ CCCTGTCTGTCTGCTACGTA 3′), UGT8 (5′ GTGGACCCTAATGATATGTG 3′), or ST3GAL5 (5′ AACTTCCGGAACCCAAAAGG 3′) (GM3S) were cloned into pLenti6.3-V5-TOPO vector (K5315-20) from Invitrogen. 293T cells were used for lentiviral packaging by co-transfection with PMD2G and Pspax2 plasmids. MDCK cells (seeding density) seeded in 6-well plates were transfected with the lentivirus containing sgRNAs for 24 h followed by puromycin (4 μg/ml) selection. The cells were passaged up to 3 times to generate stable cell lines. Knockout efficiency validation was done by real-time PCR using the following primers: UGCG (F & R), UGT8 (F: AAGACACCAAGACAAAGCCA, R: GAATTCCCAAGACCCACTCTG) and ST3GAL5 (F: CCTTCAGTACTCAGAGCCTCAG, R: CTAAGACAACGGCAATGACACC).

**Transmission electron microscopy**

Intact plasma membrane (PM) sheets of MDCK cells stably expressing GFP-KG12V, or GFP-OPR5, or GFP-ORP8 were prepared and fixed with 4% PFA and 0.1% glutaraldehyde. Next, PM sheets were then labeled with 4.5 nm gold conjugated to anti-GFP antibody followed by negative staining with uranyl acetate. The JEOL JEM-1400 transmission EM was used to acquire images of the gold particles at 100,000X magnification. ImageJ was used to process the image and to assign the coordinates of gold particles within a 1-μm2 PM area. EM-univariate analysis was used to quantify the lateral spatial distribution of a single population of immunolabeled gold nanoparticles on intact PM sheets. Briefly, the Ripley's K-function analysis tests a null hypothesis that all points in the selected area are randomly distributed:

$$K(r) = An^{-2} \sum_{i \neq j} w_{ij} \, 1(\|x_i - x_j\| \leq r) \quad \text{(A)}$$

$$L(r) - r = \sqrt{\frac{K\,(r)}{\pi}} - r \quad \text{(B)}$$

where K(r) designates the univariate K-function for the number of gold nanoparticles (n) in an intact PM area of A; r = length scale between 1 and 240 nm with an increment of 1 nm; || · || = Euclidean distance, where the condition of ||xi − xj|| ≤ r yields an indicator function of 1(·) = 1 and the condition of ||xi − xj|| > r yields indicator function of 1(·) = 0. To accomplish an unbiased edge correction, $wij^{-1}$ characterizes the fraction of the circumference of a circle that has the center at xi and radius ||xi − xj||. K(r) is transformed into *L(r) – r* then normalized against the 99% CI estimated via Monte Carlo simulations. *L(r) – r* >1.0 or < -1.0 indicates significant clustering or dispersal at the radius r, respectively. A total of 15 PM sheets per treatment were imaged, analyzed, and pooled. Bootstrapping test with 1000 times replacement was done to assess the significance between replicated point patterns.

**Electrophysiology**

Patch clamp experiments were performed at 21–25 °C using the standard whole-cell recording configuration by using HEK EPC9 USB double patch amplifier controlled by Patchmaster software (HEKA Elektronik) as previously described. Only cells with high input resistance (>2Ω) were selected for recording; membrane potentials were corrected for a liquid junction potential of 10mV. The holding potential was set to 0mV, and currents were monitored by voltage ramps of 50 ms, spanning a range of −100 to+ 100 mV, applied at 2 s intervals over a period of 100–400 s. Currents were filtered at 2.9 kHz and digitized at a rate of 20 kHz. Currents obtained before the activation of CRAC channels were assigned as leak currents and subtracted from the subsequent recorded currents. The current densities accurately measured by correcting the leak currents collected in $Ca^{2+}$ Ringer's solution with 50 μM $LaCl_3$. All patch clamp data analysis and curve fitting were done with Igor Pro 5.03 (WaveMetrics) and Graphpad prism softwares. The standard extracellular Ringer's solution contained (in mM) 120 NaCl, 2 $MgCl_2$ 10 TEACL, 10 HEPES, 10 $CaCl_2$, and 10 D-glucose (pH adjusted to 7.4 using NaOH). The standard intracellular solution contained (in mM) 120 Cesium-glutamate, 8 $MgCl_2$, 10 BAPTA, and 10 HEPES (pH adjusted to 7.2 using CsOH).

**Acknowledgements:** This work was supported by a NIH grant (R01 GM151280) and a CPRIT grant (RP250213), awarded to JL and JFH, a NIH grant (R01GM144986) and a Welch Foundation grant (BE-1913-20220331) awarded to YZ, and a NIH grant (K01HL143111) awarded to TIM. Fluorescence microscopy was performed at the Center for Advanced Microscopy, a Nikon Center of Excellence, at McGovern Medical School, UTHealth Houston (RRID: SCR_025962).
